# PKM ζ-PKC ι / λ double-knockout demonstrates atypical PKC is crucial for the persistence of hippocampal LTP and spatial memory

**DOI:** 10.1101/2025.11.10.687514

**Authors:** Panayiotis Tsokas, Changchi Hsieh, Alejandro Grau-Perales, Andrew Tcherepanov, Leo Kwok, Laura Melissa Rodriguez-Valencia, David A. Cano, Kim D. Allen, Hannah J. H. Smith, Sabina Kubayeva, Benson J. Wei, Samuel Sabzanov, Rafael E. Flores-Obando, Sourav Ghosh, Peter John Bergold, Jerry Rudy, James E. Cottrell, André Antonio Fenton, Todd Charlton Sacktor

## Abstract

PKMζ, a persistently active atypical PKC (aPKC) isoform, is thought to maintain late-phase long-term potentiation (late-LTP) and long-term memory. However, PKMζ-knockout mice still exhibit hippocampal LTP and spatial memory while lacking neocortical LTP, questioning whether this kinase is fundamental to enduring synaptic potentiation and memory. Tsokas et al. (2016) showed the other aPKC, PKCι/λ, likely compensates for PKMζ during maintenance in the hippocampus of PKMζ-null mice. In wild-type mice, PKCι/λ drives early-LTP and short-term memory, while PKMζ compensates for PKCι/λ knockout by supporting both early- and late-phase processes. Here we show PKCι/λ persistently increases during maintenance in two mouse models: PKMζ-conditional knockout (cKO) mice, and double-knockout mice carrying both conditional deletion of PKCι/λ and constitutive loss of PKMζ. In the double-knockout mice, PKCι/λ was measured while the kinase was still present, prior to its inducible ablation, to characterize its compensatory upregulation in late-LTP before removal. To test whether this compensation was functional, we ablated PKCι/λ in the hippocampus of the double-knockout mice. The double-knockout eliminated late-LTP, whereas individual knockout of either aPKC alone showed normal-appearing LTP. Double-knockout also abolished spatial long-term memory without affecting short-term memory. Thus, when PKMζ is absent, PKCι/λ persists to maintain hippocampal late-LTP and long-term memory.

## Introduction

In PKM ζ-knockout mice (PKM ζ-KO) the key long-term processes of hippocampal LTP and long-term memory appear normal, and LTP is still disrupted by the aPKC inhibitor ZIP (Lee et al., 2013; Tsokas et al., 2016; Volk et al., 2013). In the same knockout mice, LTP is eliminated in medial prefrontal cortex (mPFC) (Kniffin et al., 2025; Sacktor, 2026). These results seem inconsistent, suggesting that the persistently active PKM ζ is crucial only for LTP maintenance in mPFC, but not in hippocampus. An alternative hypothesis, however, is that compensation by another ZIP-sensitive maintenance molecule is induced in hippocampus by the absence of *Prkcz,* the gene for PKM ζ. In Tsokas et al., 2016, multiple members of the PKC gene family were found to increase expression in compensation for the loss of PKM ζ, and the most promising candidate was the other aPKC, the ZIP-sensitive PKC ι / λ, since closely related genes can compensate for one another (Conant and Wagner, 2004; El-Brolosy and Stainier, 2017; Gu et al., 2003; White et al., 2013). Notably, PKC ι / λ (referred to as PKC ι) is critical to the initial generation of the early phase of LTP and short-term memory in wild-type (WT) mice, and PKM ζ compensates for the conditional knockout of the PKCι gene (*Prkci*) to support both short- and long-term processes (Ren et al., 2013; Sheng et al., 2017; Tsokas et al., 2016; Wang et al., 2016). Perhaps LTP and long-term memory maintenance might require a persistent kinase after all — but not always PKM ζ.

The notion that the persistence of PKC ι is a biochemical mechanism responsible for late-phase LTP and long-term memory in PKM ζ -knockout mice raises two questions: 1) Does PKC ι persist in LTP and long-term memory in the absence of PKM ζ, and 2) Does eliminating both aPKCs abolish enduring LTP and long-term memory? We address these questions using conditional and double-knockout transgenic mouse strategies.

## Results

As both inducible and constitutive PKM ζ-KO mice still show hippocampal LTP (Volk et al., 2013), it is important to know if compensatory increases in other PKCs are present in the hippocampus of conditional PKM ζ-KO mice ( ζ -cKO), as was observed in PKM ζ-null mice (Tsokas et al., 2016). In PKM ζ-null mouse hippocampus, basal levels of PKC ι and the conventional PKC βI increased (Tsokas et al., 2016). To examine the hippocampus of ζ-cKO mice, adult *Prkcz*-floxed mice expressing tamoxifen-inducible Cre recombinase under the CaMKIIα-promoter (*Camk2a-CreER^T2^; Prkcz^fl/fl^* mice) were injected with tamoxifen. One week later, expression of PKM ζ in hippocampus decreased, and there was a compensatory increase in PKC ι, as well as PKC ι phosphorylated on its activation-loop (Figure 1A, Figure 1 — figure supplement 1A, Figure 1 — table supplement 1A and 2A). In addition, the expression of all four conventional PKCs (α, βI, βII, γ) increased, as did phosphorylation of the conventional PKC activation-loop (Figure 1B, Figure 1 — figure supplement 1A and 2, table supplement 1B and 2A). In contrast, there was no change in the expression of the four novel PKC isoforms ( δ, ε, η, θ) or phosphorylation of the activation-loop of PKC ε (Figure 1B, Figure 1 — figure supplement 1A and 2, table supplement 1B and 2A). There also was no change in either the level of CaMKIIα or CaMKIIα T286-autophosphorylation, which initiates Ca^2+^-independent autonomous kinase activity (Miller and Kennedy, 1986) and LTP induction (Bayer and Giese, 2024; Tullis et al., 2023) (Figure 1 — figure supplement 1B, table supplement 2B). Thus, both conditional and constitutive PKM ζ-KO mice express compensatory increases in PKC ι, as well as other PKCs, in hippocampus.

**Figure 1.**
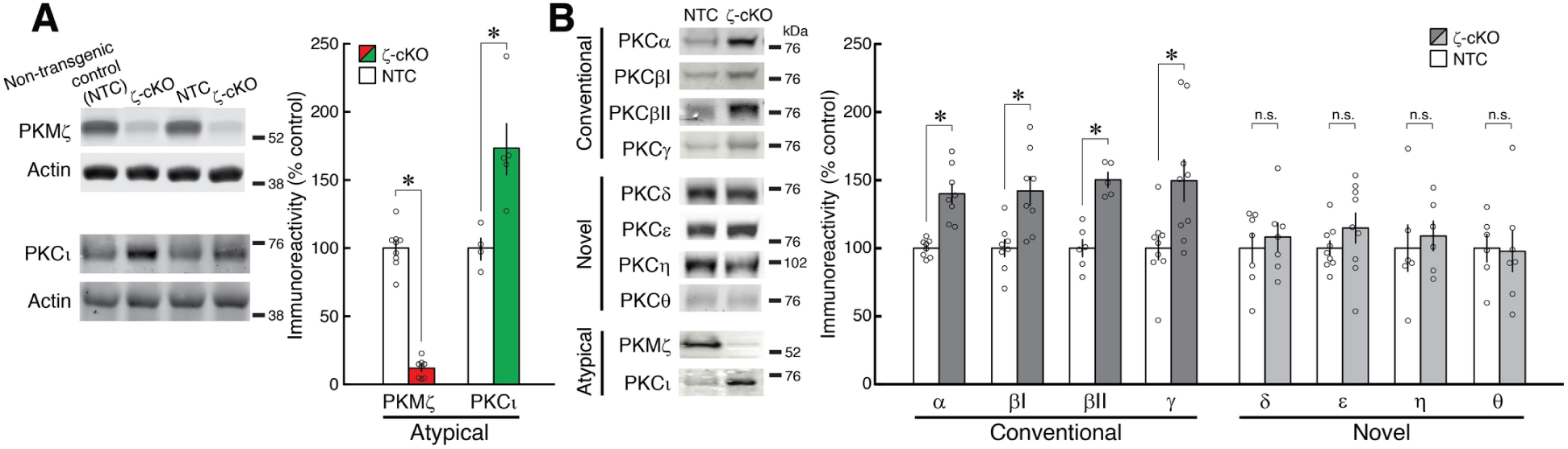
Compensatory increases of atypical PKC ι and conventional, but not novel PKCs, in conditional PKM ζ-KO mouse hippocampus. (**A, B**) Immunoblots of hippocampal extracts from *Camk2a-CreER^T2^; Prkcz^fl/fl^* mice that received tamoxifen (2 mg/200 µl i.p., 5 daily doses) to activate Cre recombinase selectively in excitatory neurons. Mice were sacrificed 7 days after the last dose. Left, representative immunoblots with Mr markers. Right, mean ± SEM. Significance by two sample Student *t* tests with Bonferroni correction denoted by *; not significant, n.s.; statistics in Figure 1 — table supplement 1A and B. Tamoxifen is a partial PKC antagonist and may still be present after a week (O’Brian et al., 1985); therefore, WT mice that also received tamoxifen are non-transgenic controls (NTC). (**A**) PKM ζ decreases and PKC ι increases in PKM ζ-cKO mice. (**B**) Conventional PKCs increase and novel PKCs do not change. Actin loading controls shown in Figure 1 — figure supplement 2.

Does the increase in PKC ι persist in long-term memory, such that one aPKC substitutes for the other? Previous research has shown that PKM ζ expression in CA1 *stratum (st.) radiatum* remains elevated for at least a month after training transgenic mice on a spatial memory task, as a component of a PKM ζ-engram that traces the tri-synaptic circuit in the dendritic compartments of memory-tagged neurons (Han et al., 2026; Hsieh et al., 2021). We determined if PKM ζ-cKO mice would also exhibit a compensatory increase of PKC ι in *st. radiatum*. The PKM ζ gene was ablated in adult *Camk2a-CreER^T2^; Prkcz^fl/fl^* mice (Figure 2A), and 3 weeks later the mice were trained on the active place avoidance task to produce spatial memory. One week after training, compared to vehicle-injected littermates, the PKMζ -cKO mice showed a decrease to ∼25% in PKM ζ expression and an increase to ∼400% in PKC ι expression (Figure 2B, Figure 2 — table supplement 1), along with spatial memory retention (Figure 2C). Thus, spatial conditioning of ζ-cKO mice creates long-term active place avoidance memory and induces compensatory, persistent increased expression of hippocampal PKC ι.

**Figure 2.**
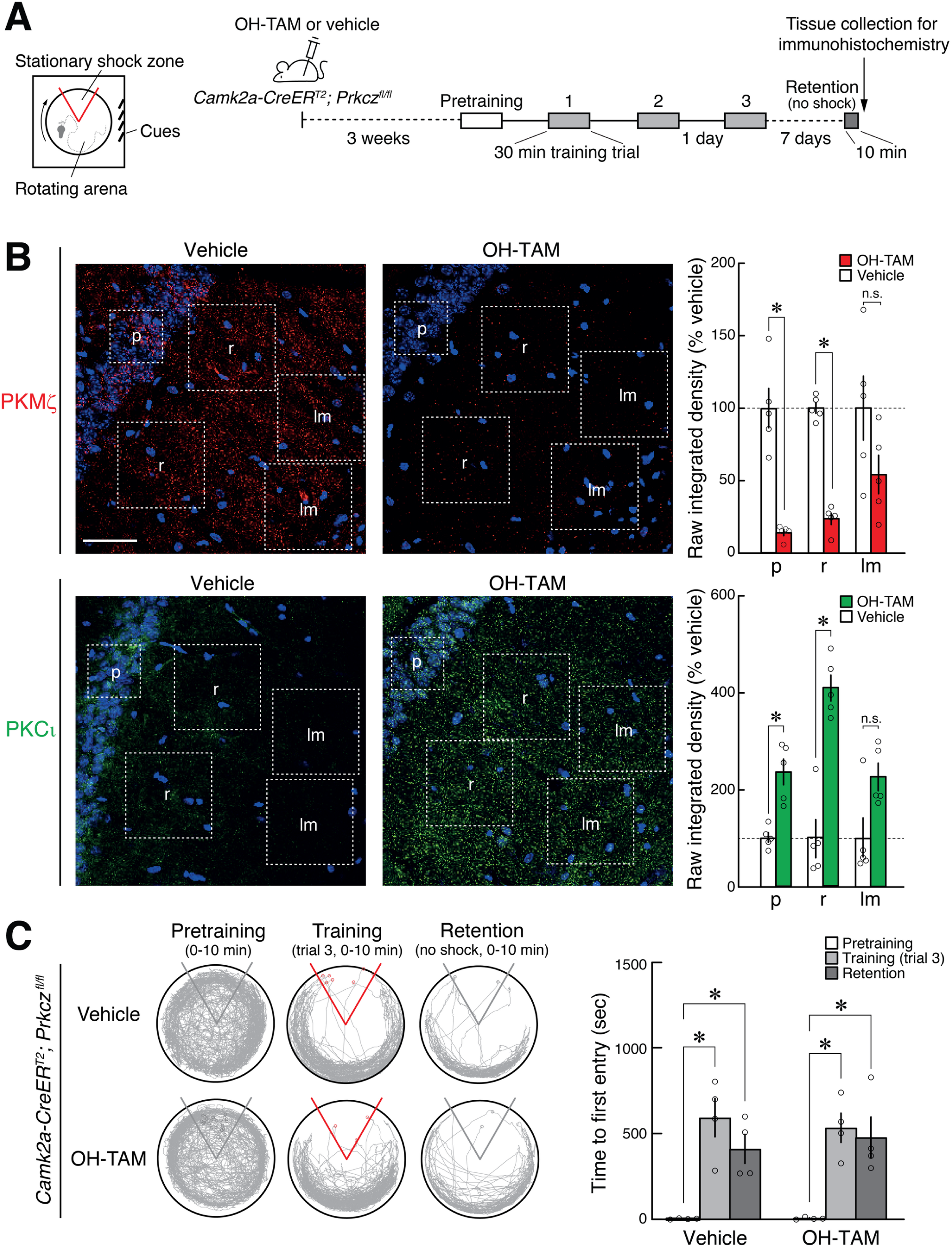
Compensatory increases of PKC ι during spatial memory in conditional PKM ζ-KO mouse hippocampus. (**A**) Left, schematic of active place avoidance training apparatus with a slowly rotating arena containing a nonrotating shock zone sector (shown in red). Visual cues located on the walls of the room are needed to avoid the shock zone. Right, experimental protocol. PKM ζ is genetically ablated in *Camk2a-CreER^T2^; Prkcz^fl/fl^* mice. Cre is activated using 4-OH tamoxifen (OH-TAM, 2 mg/200 µl i.p., 3 doses every other day). Control mice receive vehicle injections. Active place avoidance training begins 3 weeks later, and 1 week after training memory retention is tested in the absence of shock followed by sacrifice and immunohistochemistry. (**B**) Immunohistochemistry shows ζ-cKO reduces PKM ζ and increases PKC ι in CA1 *st. pyramidale* (p) and *radiatum* (r), but not *lacunosum-moleculare* (lm) 1 week after training. Left above, PKM ζ expression decreases in cell bodies and dendritic compartments of the PKM ζ-cKO. Left below, PKC ι expression increases in cell bodies as well as in dendritic compartments where it is ordinarily expressed at low levels. DAPI staining of nuclei shown in blue. Bar = 50 µm. Right, mean ± SEM. Student *t* tests with Bonferroni corrections compared differences in PKM ζ and PKC ι expression separately in the CA1 *strata* (Figure 2 — table supplement 1). (**C)** Compensatory spatial memory in ζ -cKO. Left, representative paths during first 10-min of pretraining, training trial 3, and 1-day memory retention. Right, mean ± SEM. Two-way ANOVA (treatment X training) revealed a significant effect of training (*F*_1.662, 9.972_ = 41.93, *P* < 0.0001), but not an effect of treatment (*F*_1, 6_ < 0.001, *P* = 1.0) or their interaction (*F*_2, 12_ = 0.48, *P* = 0.6). Further comparisons using Bonferroni-corrected tests revealed significant differences between pretraining and training (Trial 3) (*P* = 0.002) and pretraining and retention (*P* = 0.001), but no differences between training (Trial 3) and retention (*P* = 0.1). Further comparison revealed significant differences between pretraining and training (Trial 3) in both vehicle and 4-OH tamoxifen groups (*P* = 0.02 and *P* = 0.01, respectively) and between pretraining and retention in both vehicle and 4-OH tamoxifen groups (*P* = 0.03 and *P* = 0.05, respectively), confirming the treatment groups did not behave differently.

Does the persistent PKCι expression substitute for the function of PKMζ in LTP and memory of PKM ζ-KO mice? As PKC ι -null mice are embryonically lethal (Seidl et al., 2013), we determined the functional significance of the compensatory increase of PKC ι for LTP by injecting an adeno-associated virus (AAV) expressing Cre-recombinase in one hippocampus of PKC ι -floxed/PKM ζ-null (*Prkci^fl/fl^; Prkcz^−/–^*) mice to produce a double-knockout (dKO) (Sheng et al., 2017) (Figure 3A). The contralateral hippocampus was injected with a control AAV expressing eGFP, and 3 weeks later *ex vivo* slices were prepared. PKC ι decreased in the ipsilateral hippocampus to ∼20% compared to the contralateral hippocampus (Figures 3A and B).

**Figure 3.**
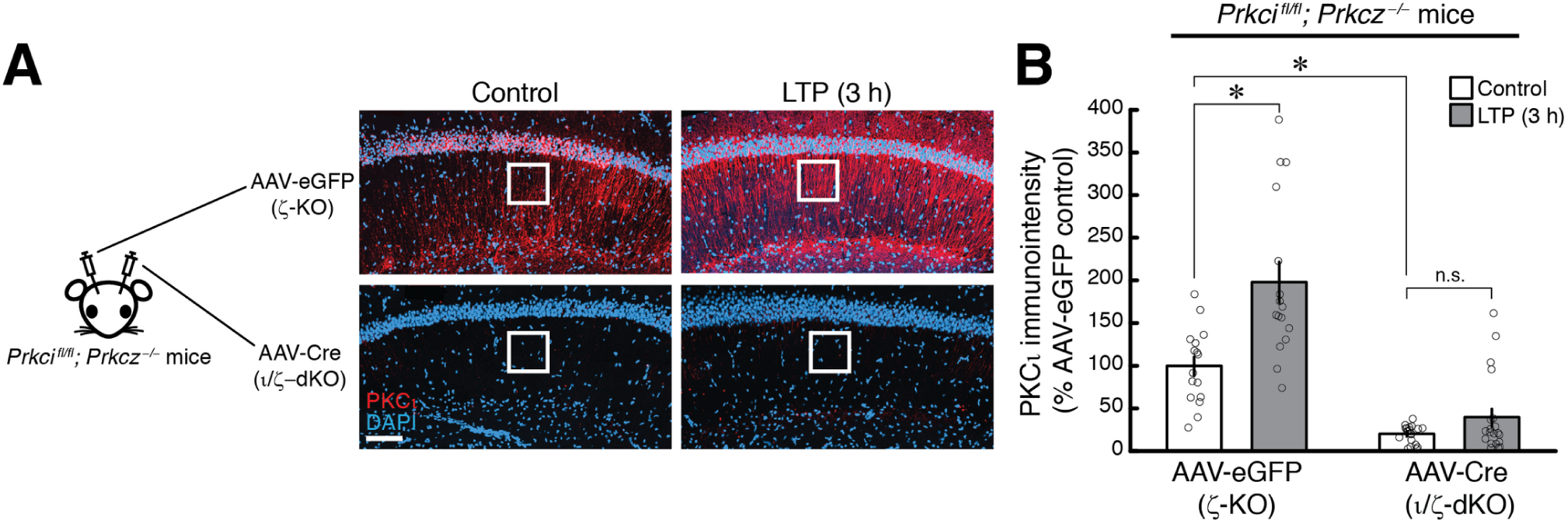
Compensatory increases of PKC ι during hippocampal late-LTP maintenance in *Prkci^fl/fl^; Prkcz^−/–^* mice. (**A**) Left, schematic of experimental protocol shows AAV expressing Cre by cytomegalovirus (CMV) promoter injected into ipsilateral hippocampus of a *Prkci^fl/fl^; Prkcz^−/–^*mouse, and control AAV expressing eGFP injected into contralateral hippocampus. Hippocampal slices are prepared 3 weeks later. Right, representative images of PKC ι - immunohistochemistry from adjacent slices in AAV-eGFP-injected ( ζ-KO) hippocampus show PKC ι persistently increases 3 h post-tetanization (top row), and low, unchanging levels of PKC ι in the AAV-Cre-injected ( ι / ζ -dKO) hippocampus (bottom row). White boxes show *st. radiatum* regions of interest. DAPI staining of nuclei shown in blue. (**B**) Mean ± SEM. The two-way ANOVA reveals the main effects of treatment (AAV-Cre [ ι / ζ-dKO] vs. AAV-eGFP [ ζ-KO], *F*_1,68_ = 83.58, *P* < 0.00001, *η*^2^_p_ = 0.55), and stimulation (HFS vs. test, *F*_1,68_ = 20.47, *P* = 0.00003, *η*^2^_p_ = 0.23), and an interaction of treatment X stimulation (*F*_1,68_ = 9.09, *P* = 0.004, *η*^2^_p_ = 0.12). *Post-hoc* analysis confirms that, compared to the ζ-KO control group, the intensity of PKC ι immunoreactivity was significantly decreased in ι / ζ -dKO (*P*’s < 0.002 for both control and LTP in ι / ζ -dKO), and increased in ζ-KO after HFS (*P* = 0.00011, ζ-KO, n’s = 16, ι / ζ- dKO, n’s = 20). Intensity of PKC ι immunoreactivity did not change in the ι / ζ-dKO between the control and HFS groups (*P* = 0.3). Bar = 100 µm.

If PKC ι is important for enduring LTP in PKMζ -KO mice, then this decrease should disrupt LTP. High-frequency stimulation (HFS) of Schaffer collateral/commissural-CA1 synapses in the ipsilateral ι / ζ-dKO slices produced no persistent change in the residual PKC ι and a transient LTP only lasting ∼1-2 h (Figures 3 and 4A; Figure 4 — figure supplement 1A). In contrast, HFS of the contralateral slices, expressing PKC ι but not PKM ζ, induced persistent increases of the PKC ι to ∼200% and compensatory late-LTP, both lasting at least 3 h, the duration of the recordings (Figures 3 and 4A, Figure 4 — figure supplement 1A). Slices from hippocampus of *Prkci^fl/fl^; Prkcz^−/–^* mice injected with AAV expressing Cre by the CaMKIIα-promoter to selectively ablate the PKC ι gene in excitatory neurons resulted in similar transient LTP (Figure 4A, inset; Figure 4 — figure supplement 1B and 2A). In addition, we showed the virus injected into PKCι - floxed mice that express PKM ζ (*Prkci^fl/fl^; Prkc^+/+^* mice) resulted in compensated early-LTP, as previously reported (Sheng et al., 2017) (Figure 4 — figure supplement 1C and 2B). Thus, individual knockout of each aPKC shows compensatory LTP, whereas double-knockout of both aPKCs eliminates late-LTP.

**Figure 4.**
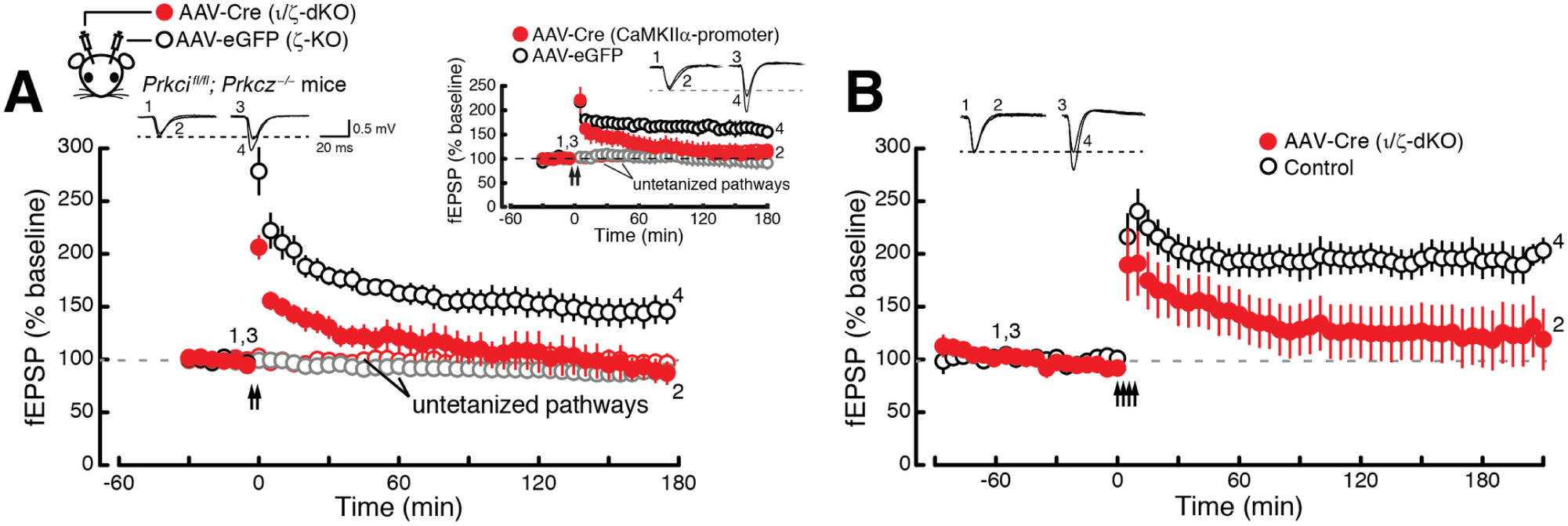
ι / ζ-dKO hippocampus shows transient LTP, but not persistent LTP. (**A**) Late-LTP is absent in ι / ζ-dKO hippocampus. Above left inset, schematic of intrahippocampal injections of AAC-Cre recombinase and AAV-eGFP. Middle inset, representative fEPSPs correspond to numbered times in time-course below. Below, filled red circles, AAV expressing Cre and HFS with 2 tetanic trains; open red circles, test stimulation of a second synaptic pathway within the hippocampal slice. HFS tetani shown at arrows. Open black circles, AAV expressing eGFP by CMV promoter with HFS; open grey circles, with test stimulation. Three-way mixed-design ANOVA reveals main effects of treatment (hippocampal injections of AAV-Cre [ ι / ζ-dKO] vs. AAV-eGFP [ζ-KO], *F*_1,20_ = 8.45, *P* = 0.0009, *η*^2^_p_ = 0.30), and stimulation (HFS vs. test stimulation, *F*_1,20_ = 5.90, *P* = 0.025, *η*^2^_p_ = 0.23), as well as a 3-way interaction among treatment X stimulation X time (5-min average of pre-HFS and 3-h post-HFS, *F*_1,20_ = 12.68, *P* = 0.002, *η*^2^_p_ = 0.39). *Post-hoc* analysis confirms established LTP is not maintained in ι / ζ-dKO 3 h after HFS when compared to pre-HFS basal responses (*P* = 0.7). *Post-hoc* analysis also confirms the control hippocampus maintains established LTP (*P* = 0.0002). Test stimulation was unaffected by AAV-Cre or AAV-eGFP injections (*P* = 0.9 and *P* = 0.7, respectively, n’s = 6). Right inset, ι / ζ -dKO by CaMKIIα promoter expression of Cre eliminates late-LTP. Three-way mixed-designed ANOVA reveals interaction between treatment (ζ-KO vs. ι / ζ -dKO) and stimulation (HFS vs. test stimulation, *F*_1,14_ = 6.62, *P* = 0.02, *η*^2^_p_ = 0.32), and a 3-way interaction among treatment, stimulation, and time (5 min pre-HFS and 3 h post-HFS, *F*_1,14_ = 8.56, *P* = 0.01, *η*^2^_p_ = 0.38). *Post-hoc* analysis confirms that compared to pre-HFS basal responses, LTP is not maintained in ι / ζ-dKO hippocampus 3 h post-HFS (*P* = 0.8) and is maintained in the control hippocampus (*P* = 0.003). Test stimulation was unaffected by AAV-Cre or AAV-eGFP injections (*P* = 0.4 and *P* = 0.9, respectively). ι / ζ -dKO HFS, n = 5; ι / ζ -dKO test, n = 4; ζ-KO HFS, n = 5; ζ-KO test, n = 4. (**B**) LTP does not persist in ι / ζ-dKO mice after stronger afferent stimulation with 4 tetanic trains. ANOVA with repeated measurements reveals main effects of time (5 min pre-HFS, 20 min post-HFS, and 3 h post-HFS, *F*_2,14_ = 20.51, *P* < 0.0001, *η*^2^_p_ = 0.75). *Post-hoc* analysis confirms that early-LTP is established in both ι / ζ-dKO and control groups (5 min pre-HFS vs. 20 min post-HFS, *P* = 0.005 and 0.002, respectively), and no difference between these two groups at 20 min post-HFS (*P* = 0.6). However, LTP in ι / ζ-dKO did not persist 3 h (5 min pre-HFS vs. 3 h post-HFS, *P* = 0.4), whereas LTP is intact in control. (*P* = 0.008). ι / ζ -dKO, n = 5; control, n = 4.

We tested if late-LTP could be induced in ι / ζ-dKO hippocampus by increasing HFS from two trains, 20 sec apart, which is optimized to produce an early onset of late-LTP (Tsokas et al., 2007), to four trains, spaced 5 min apart, which is optimized to produce maximal late-LTP (Scharf et al., 2002; Serrano et al., 2005) (Figure 4B; Figure 4 — figure supplement 1D). The stronger stimulation induces LTP in ι / ζ -dKO slices that lasts only ∼1-2 h. This LTP was no longer expressed at 3 h post-stimulation, at which time field excitatory postsynaptic potentials (fEPSPs) did not significantly differ from baseline fEPSPs before HFS.

To determine if PKC ι supports long-term memory in the absence of PKM ζ, we compared the effects on spatial memory of ι / ζ- dKO to ζ-KO by injecting *PKC ι ^fl/fl^; PKM ζ^−/–^*littermates bilaterally in hippocampus with either AAV-Cre or AAV-eGFP (Figure 5A). Mice received three 30-min training trials separated by 24 hours and a final retention test without shock the next day. We assessed two measures of short-term memory (Figure 5B). First, we examined the time to each entry into the shock zone in the first training trial, as compared to the entries into the shock zone during the pretraining session with the shock off. This avoidance behavior increased within the first 30-min training session in both the ι / ζ -dKO and ζ -KO mice, and the increases were indistinguishable. Second, we measured the maximum avoidance time within each session. Maximum avoidance time reflects the time between shocks, which is controlled by the animal’s behavior and within-trial memory. Note that compared to pretraining between-shock zone entries, the maximum avoidance times for ζ-KOs and ι / ζ-dKOs increased in the first training trial, indicating both genotypes acquired short-term memory for the shock zone. The increases in the two groups were indistinguishable. The maximum avoidance time for the ζ-KO, however, increased over the 3 daily training sessions, whereas that of the ι / ζ -dKO did not, suggesting impaired long-term memory in the ι / ζ- dKO compared to the ζ -KO. Our main measure of long-term memory was time to first entry into the shock zone at the beginning of each session (Figure 5C). This increases with long-term memory maintained across days from previous trials. The ζ-KOs’ first entry times increased dramatically from pretraining to both session 3 and retention test, indicating that mice with PKC ι maintain long-term memory. In contrast, ι / ζ -dKO mice displayed a minimal increase at session 3 that was not significantly different from pretraining, and no detectable difference between pretraining and the retention test, indicating the loss of spatial long-term memory.

**Figure 5.**
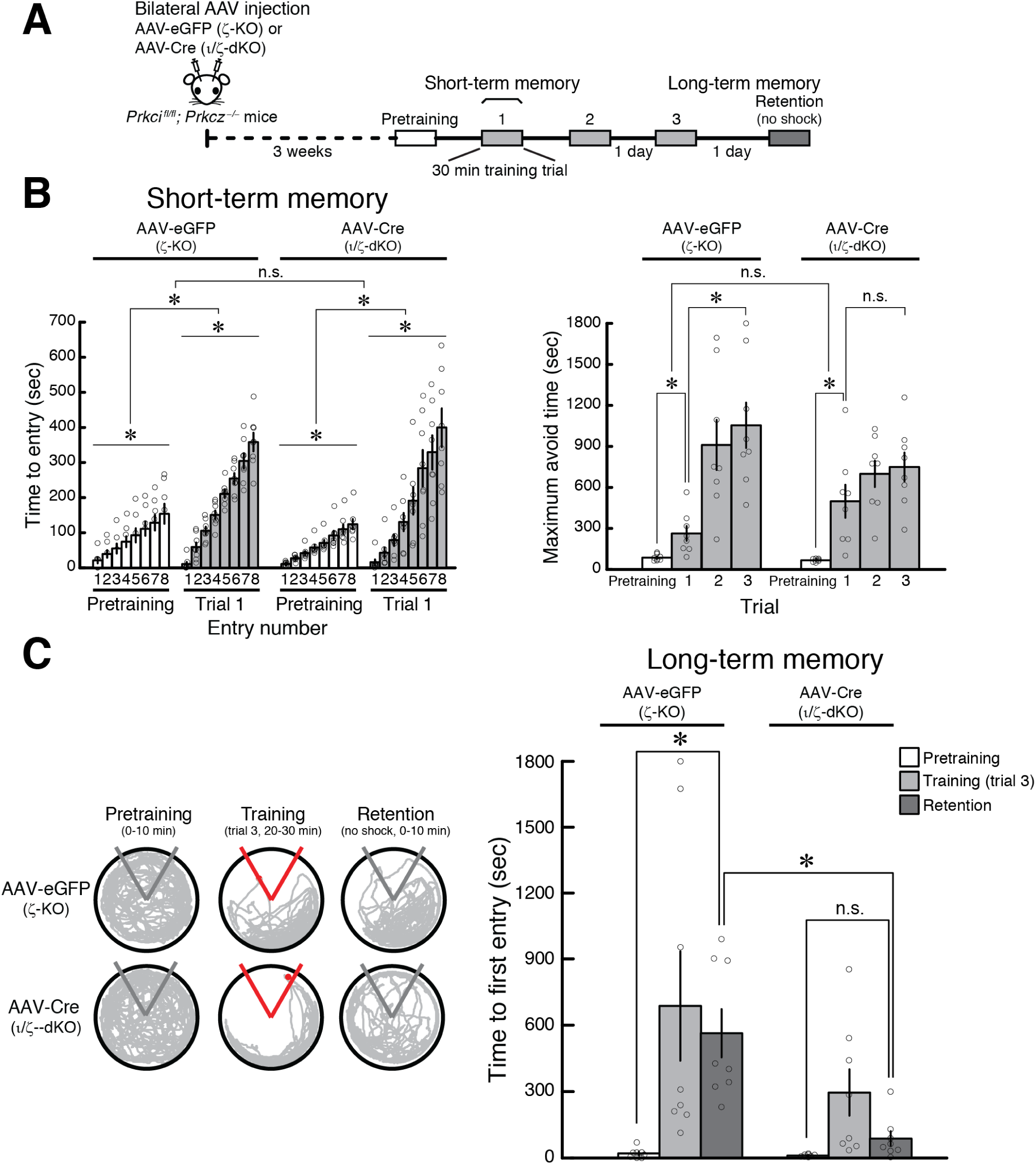
Impaired long-term memory and intact short-term memory for spatial information in mice with bilateral hippocampal ι / ζ-dKO. (**A**) Experimental protocol. *Prkci^fl/fl^; Prkcz^−/–^* mice are injected bilaterally in hippocampus with AAV-Cre ( ι / ζ-dKO) or AAV-eGFP ( ζ-KO, control), and 3 weeks later they received pretraining and, after 1 day, a single 30 min trial repeated daily for a total of 3 trials. Long-term retention is tested without shock 1 day after the last training trial. (**B**) ι / ζ-dKO does not affect short-term memory in the first training trial. Left, ι / ζ -dKO does not affect short-term memory as assessed by the time to enter the shock zone for the first 8 times (all animals had up to at least 8 entries in trial 1). ANOVA with repeated measurements finds the main effects of training (pretraining and trial 1, *F*_1,28_ = 35.19, *P* < 0.00001, *η*^2^_p_ = 0.56) indicating trial 1 learning, time to entry (the 1^st^ to 8^th^ entry within a trial, *F*_7,196_ = 145.80, *P* < 0.00001, *η*^2^_p_ = 0.84), and their interaction (*F*_7,196_ = 37.68, *P* < 0.00001, *η*^2^_p_ = 0.57). However, there is no group effect (AAV-eGFP- and AAV-Cre-injected, *F*_1,28_ = 0.19, *P* = 0.7, *η*^2^_p_ = 0.007) nor interaction with either training or time to each entry (*F*’s < 0.66, *P*’s> 0.60, *η*^2^_p_’s < 0.02). Right, ι / ζ-dKO does not affect short-term memory as assessed by maximum avoidance time during the first training trial. The contrast analysis reveals that the increases of maximum avoidance time from pretraining to trial 1 are not different between AAV-eGFP-injected and AAV-Cre-injected groups (*t*_14_ = 1.91, *P* = 0.08, *d* = 1.91). Paired *t*-tests reveal trial 1 is greater than pretraining in each genotype (*t*’s > 3.10, *P*’s < 0.018, Cohen’s *d*’s > 1.62), indicating both groups of mice successfully established short-term memory. In contrast, the improvement of maximum avoidance time from trial 1 to trial 3 are different between the groups (*t*_14_ = 2.93, *P* = 0.01, *d* = 2.88), suggesting the two groups performed differently between daily training sessions when between-day memory influences avoidance. In addition, the ANOVA with repeated measurement discovers no group effect (AAV-eGFP-injected vs. AAV-Cre-injected, *F*_1,14_ = 0.56, *P* = 0.47, *η*^2^_p_ = 0.04), but significant effects of trial (*F*_3,42_ = 30.37, *P* < 0.0001, *η*^2^_p_ = 0.68), and interaction (*F*_3,42_ = 2.93, *P* = 0.04, *η*^2^_p_ = 0.17). *Post-hoc* tests confirm that the maximum avoidance time in trial 1 is not different between the two groups (*P* = 0.14). The AAV-eGFP-injected group improved their performance between trial 1 and trial 3 (*P* = 0.0002), whereas the AAV-Cre showed no improvement (*P* = 0.2; n’s = 8). These data indicate no differences in short-term memory between AAV-eGFP- and AAV-Cre-injected groups, but only the AAV-Cre-injected group failed to improve between daily trials, suggesting inability to retain avoidance memory across days. (**C**) PKCι gene ablation impairs long-term memory in *Prkci^fl/fl^; Prkcz^−/–^* mice. Left, representative paths during 10-min of pretraining, at end of training trial 3, and 1-day memory retention. Right, mean ± SEM. The ANOVA with repeated measurement finds main effects of group (AAV-eGFP vs. AAV-Cre, *F*_1,14_ = 10.53, *P* = 0.006, *η*^2^_p_ = 0.43) and training phase (pretraining, trial 3 of training, retention, *F*_2,28_ = 7.65, *P* = 0.002, *η*^2^_p_ = 0.35). *Post-hoc* analysis reveals that the mice with AAV-Cre-injected ι / ζ-dKO hippocampus perform poorer during the memory retention test, compared to AAV-eGFP-injected littermates (*P* = 0.02). The mice with ι / ζ-dKO hippocampus show no difference between the memory retention test and pretraining trial (*P* = 0.9), whereas the AAV-eGFP-injected mice show long-term memory is maintained (*P* = 0.02; n’s = 8). In addition, pretraining vs. training trial 3 was significantly different in ζ -KO (*P* = 0.006), but not in ι / ζ-dKO (*P* = 0.4).

## Discussion

Here we found that a second aPKC becomes persistently active to maintain late-LTP and long-term memory in the PKM ζ-κΟ. T he isoform most closely related to PKM ζ, PKC ι, which normally plays only a transient role in LTP and short-term memory in WT mice, persistently increases expression in LTP and long-term memory in PKMζ -KO mice (Figures 2 and 3). Although LTP is present if PKM ζ or PKC ι is individually knocked out, when both are genetically eliminated by double-knockout there is no enduring hippocampal LTP or long-term spatial memory (Figures 4 and 5, Figure 4 — figure supplement 2B).

The ι / ζ- dKO exhibits an early transient synaptic potentiation that could not be maintained into a late-phase of LTP (Figure 4, Figure 4 — figure supplement 1A, B, and D). In addition, bilateral hippocampal ι / ζ -dKO eliminated long-term spatial memory but did not prevent learning, short-term memory, or expression of the place avoidance behavior (Figure 5). As PKC ι is a key contributor to early-LTP and short-term memory in WT mice (Ren et al., 2013; Wang et al., 2016), there must be additional compensation for short-term processes present in the aPKC-dKO.

Our finding that PKC ι can substitute for PKM ζ raises the question of how the compensation is induced. After gene knockout, compensatory gene expression can be triggered by the fragments of mRNA that are produced by transcription upstream of the site of Cre-recombination (El-Brolosy et al., 2019; El-Brolosy and Stainier, 2017; Ma et al., 2019). This could explain why constitutive and conditional PKM ζ-KOs produce compensation and normal-appearing late-LTP/long-term memory, whereas PKM ζ-shRNA and PKM ζ-antisense oligodeoxynucleotides that retain full-length PKM ζ mRNA transcription in WT mice do not induce compensation and disrupt late-LTP/long-term memory (Tsokas et al., 2016; Wang et al., 2016). In contrast to hippocampus, both early- and late-LTP are eliminated in prefrontal cortex of PKM ζ-KO mice (Kniffin et al., 2025). This suggests that the PKC ι activation driving early-LTP in the hippocampus of WT mice may not be available to compensate for the loss of PKM ζ in the prefrontal cortex of ζ-KO mice (Sacktor, 2026).

Once increased, how does PKC ι accomplish maintenance? PKM ζ maintenance is linked to the kinase’s second messenger-independent, persistent enzymatic activity (Sacktor et al., 1993). PKM ζ is autonomously active because the kinase is an independent catalytic domain that lacks the PKC ζ autoinhibitory regulatory domain (Hernandez et al., 2003; Sacktor et al., 1993). PKCι, however, is a full-length PKC isoform with a regulatory domain that inhibits its catalytic domain. Therefore, for it to compensate for PKM ζ, PKC ι requires additional posttranslational mechanisms to persistently activate and localize the kinase at active synapses (Figures 2B and 3). PKCι can be activated by postsynaptic proteins such as p62 that bind to its regulatory domain (Jiang et al., 2009; Ren et al., 2013). This protein-protein interaction may sustain PKCι kinase action longer than the rapidly metabolized lipid second messengers that transiently stimulate PKC ι and the conventional/novel PKCs, thus allowing one persistently active aPKC to replace the other.

The maintenance properties of PKMζ depend not only on continuous activity but also on continuous binding to the postsynaptic scaffolding protein KIBRA/WWC1 (kidney and brain protein/WW and C2 domain protein 1) (Tsokas et al., 2024). This sustained interaction perpetually targets PKMζ to active synapses in a process of persistent synaptic tagging (Hsieh et al., 2026; Shouval et al., 2025; Tsokas et al., 2024). KIBRA also binds to PKC ι, albeit more weakly than PKM ζ (Tsokas et al., 2024). Therefore, in WT mice, the strong binding of PKM ζ to KIBRA might allow it to displace PKC ι at active synapses in the transition from early- to late-LTP. In contrast, in the hippocampus of PKM ζ-KO mice, the PKCι at active synapses would not be replaced by PKM ζ. PKM ζ and PKC ι also compete for binding to PAR3 (partitioning defective protein 3), another postsynaptic protein that localizes aPKCs within neurons (Parker et al., 2013; Zhang and Wei, 2022). Thus, in the PKM ζ-KO, synaptic tags such as KIBRA and PAR3 that are components of the PKM ζ maintenance mechanism may now shift to PKC ι to sustain hippocampus-dependent LTP and long-term memory.

Our finding that persistent biochemical action by PKM ζ or PKC ι is crucial for maintaining synaptic potentiation and memory appears at odds with the widely held notion that synapses sustain memory through stable changes in structure without the need for ongoing enzymatic activity dedicated to storing information. The structural view, as introduced, hypothesized, and established by Ramón y Cajal, Hebb, and Kandel, has led to identifying structural plasticity and non-enzymatic structural molecules that contribute to establishing LTP and memory (Bailey and Kandel, 1993; Hebb, 1949; Ramón y Cajal, 1894), including cytoskeletal actin and the building blocks of perineuronal nets (Matus, 2000; Tsien, 2013). Memory maintenance by ongoing, persistent biochemical processes was an alternative hypothesis proposed by Crick, Lisman, and Schwartz (Crick, 1984; Lisman, 1985; Schwartz, 1993). The search for a biochemical process that maintains LTP for hours and long-term memory for days has focused on two persistently active protein kinases, CaMKII and PKM ζ (Lisman, 2017; Sacktor and Fenton, 2018). The kinase action of CaMKII, however, is only crucial for initiating but not perpetuating LTP and memory (Bayer and Giese, 2024; Tullis et al., 2023). By contrast, PKM ζ under the normal physiological conditions of WT mice, and PKC ι as part of the compensatory response of PKMζ -KO mice, play similar crucial roles in maintaining the molecularly distinct, late phase of LTP and long-term memory (Pastalkova et al., 2006; Shema et al., 2011; Shema et al., 2007; Tsokas et al., 2024; Wang et al., 2016). Future work will be required to determine if persistent changes in synaptic structure are sustained by the ongoing aPKC activity that maintains long-term memory (Chen et al., 2014).

## Acknowledgements

P.T. is an Alexander S. Onassis Public Benefit Foundation Scholar.

## Author contributions

Conceptualization: TCS, AAF

Methodology: PT, CH, AG-P, LK, SG

Investigation: PT, CH, AG-P, LK, LMR-V, DAC, KDA, HJHS, SK, BJW, SS, REF-O

Visualization: CH, AG-P

Funding acquisition: TCS, AAF, JEC

Project administration: TCS, AAF

Supervision: TCS, AAF

Writing – original draft: TCS, AAF, PJB, JR

Writing – review & editing: TCS, AAF, PJB, JR, JEC

## Data Availability

All data are available in the main text or the supplementary materials.

## Supplementary Materials

**Key Resources Table**

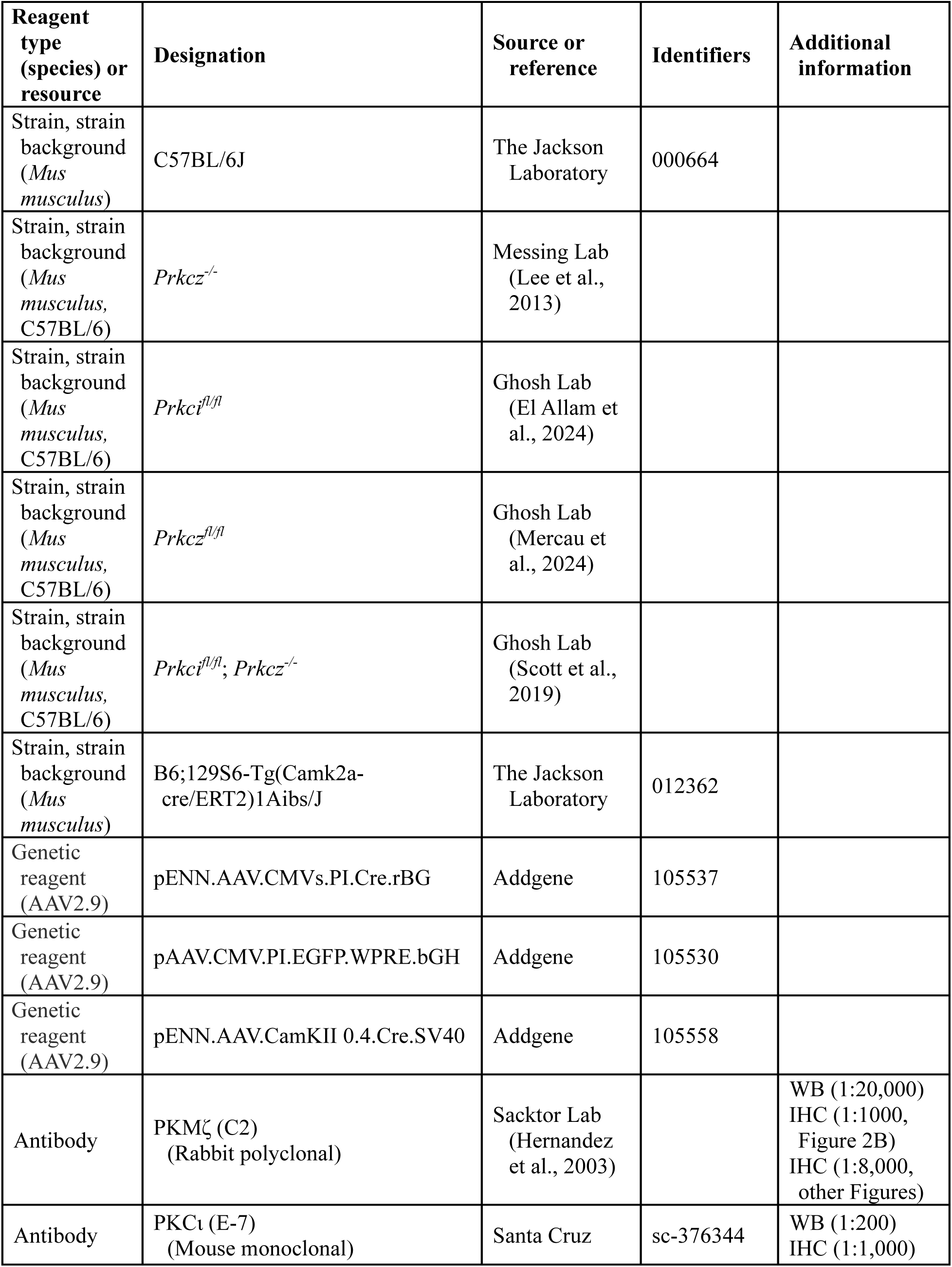
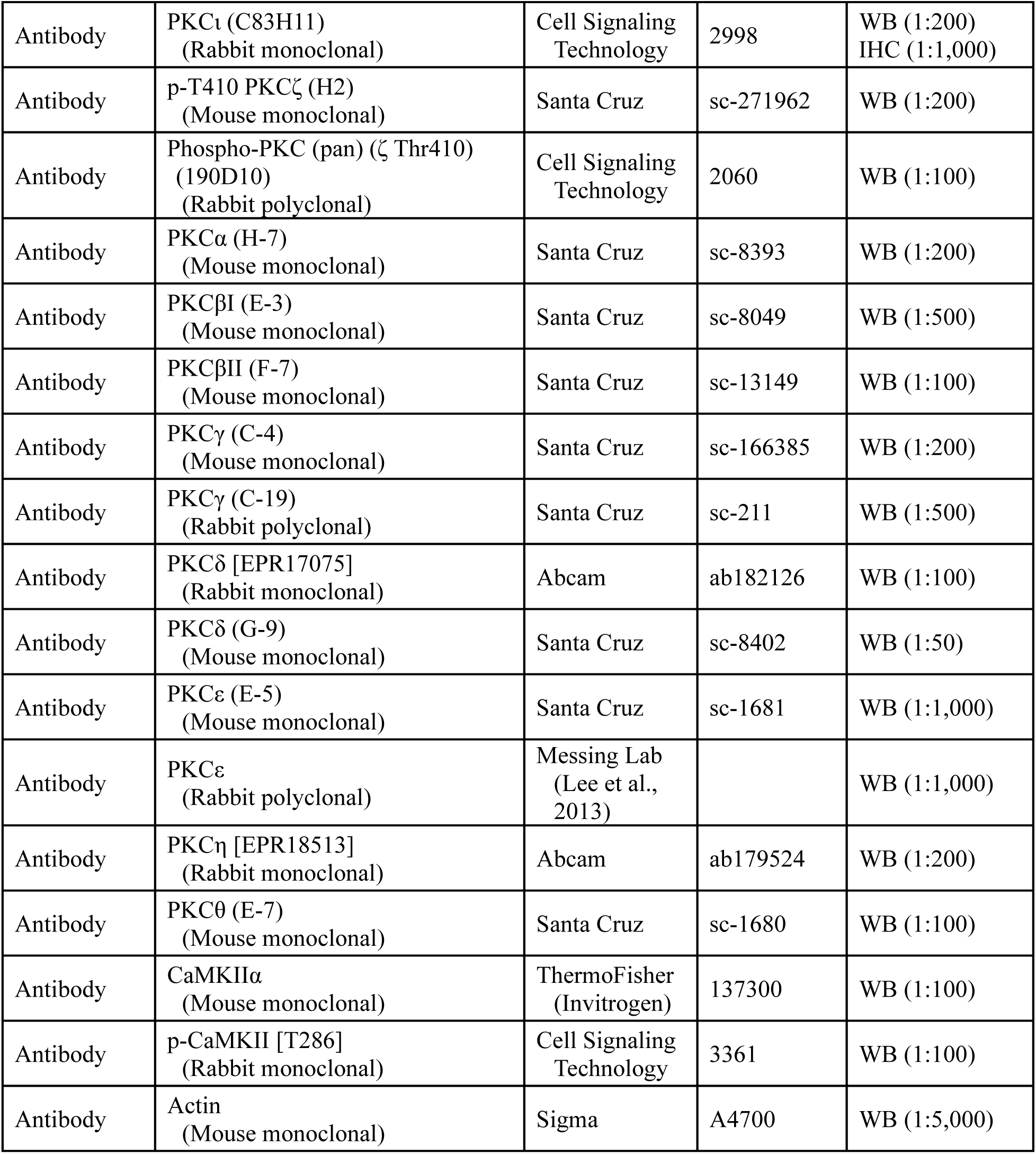

## Materials and Methods

### Reagents

Reagents were from MilliporeSigma unless otherwise stated.

### Animals

Male mice were on C57BL/6 background and at least 4-months-old for all experiments. The PKM ζ-null mouse line was previously described (Lee et al., 2013) and provided by Robert O. Messing (Univ. Texas at Austin, TX, USA). Conditional PKM ζ and PKC ι mice were generated by Sourav Ghosh as previously described (El Allam et al., 2024; Mercau et al., 2024; Scott et al., 2019). The vehicle for tamoxifen i.p. injections was sunflower seed oil for *Camk2a-CreER^T2^*mice, which were from Jackson Labs.

### Hippocampal slice recording and stimulation

Acute mouse hippocampal slices (450 µm) were prepared as previously described (Tsokas et al., 2016; Tsokas et al., 2019). Hippocampi were dissected, bathed in ice-cold dissection buffer, and sliced with a McIlwain tissue slicer in a cold room at 4°C. The dissection buffer contained (in mM): 125 NaCl, 2.5 KCl, 1.25 NaH_2_PO_4_, 26 NaHCO_3_, 11 glucose, 10 MgCl_2_, and 0.5 CaCl_2_, and was bubbled with 95% O_2_/5% CO_2_ to maintain pH at 7.4. After dissection the slices were transferred to an Oslo-type interface recording chamber (31.5 ± 1°C) (Tsokas et al., 2019). The recording superfusate consisted of (in mM): 118 NaCl, 3.5 KCl, 2.5 CaCl_2_, 1.3 MgSO_4_, 1.25 NaH_2_PO_4_, 24 NaHCO_3_, and 15 glucose, bubbled with 95% O_2_/5% CO_2_, with a flow rate of 0.5 ml/min.

Field EPSPs were recorded with a glass extracellular recording electrode (2–5 MΩ) placed in the CA1 *st. radiatum*, and concentric bipolar stimulating electrodes (CBBRE75 and 30200; FHC, Bowdoin, ME) were placed on either side within CA3 or CA1. Test stimulation rate was once every 30 sec in each stimulating electrode, alternating every 15 sec between electrodes. Based upon a pre-established exclusion criterion, a slice was not used if fEPSP spike threshold was < 2 mV on initial input-output analysis. Pathway independence was confirmed by the absence of paired-pulse facilitation between the two pathways. A single stimulating electrode with a test stimulation rate of once every 30 sec was used for immunohistochemistry experiments. HFS optimized to produce a relatively rapid onset of protein synthesis-dependent late-LTP consisted of two 100 Hz-1 s tetanic trains, at 25% of spike threshold, spaced 20 sec apart (Tsokas et al., 2005). HFS optimized to produce maximal late-LTP consisted of four 100 Hz-1 s tetanic trains, at 25% of spike threshold, spaced 5 min apart (Scharf et al., 2002; Serrano et al., 2005). The maximum slope of the rise of the fEPSP was analyzed on a PC using the WinLTP data acquisition program (Anderson and Collingridge, 2007).

### Immunoblots and Immunohistochemistry

Immunoblots of total hippocampus were performed as previously described, using antibodies in the Key Resources Table (Tsokas et al., 2016). Immunoblots were stained with multiple antisera to visualize multiple PKCs on the same immunoblot. Isoforms with similar molecular weights (e.g., the cPKCs) were either stained with antisera of different species or examined on separate blots. To conserve antisera, after transfer, the nitrocellulose was trimmed into sections containing the PKCs and actin lanes based on visible molecular weight markers.

Quantitative immunohistochemistry for Figure 3 and Figure 4 — figure supplement 2 was as described (Hsieh et al., 2021; Tsokas et al., 2024), using mouse anti-PKC ι primary antibody (1:1000, E-7, Santa Cruz SC-376344).

Quantitative immunohistochemistry for Figure 2B was performed as follows. Free-floating sections were permeabilized with phosphate-buffered saline (PBS) containing 0.1% Tween20 (PBS-T) for 1 h at room temperature and blocked with 10% normal goat serum in PBS-T (blocking buffer) for 2.5 h at room temperature. One batch of sections was incubated overnight at 4 °C with rabbit anti-PKM ζ C-2 antisera primary antibody (1:1,000) (Hernandez et al., 2003) and a second batch of sections with rabbit anti-PKC ι (1:1000, Cell Signaling #2998S) in blocking buffer. After washing 3 times for 10 min each in PBS-T, both batches of sections were incubated with the secondary antibody goat anti-rabbit-Alexa 647 (1:500 in blocking buffer; Jackson ImmunoResearch) for 2 h at room temperature. After washing 3 times for 10 min each in PBS-T and extensive washing with PBS, the sections were mounted with Vectashield with 4′,6-diamidino-2- phenylindole (DAPI, Vector Laboratories). Three sections from each mouse were examined using an upright Leica SP8 confocal microscope and analyzed using ImageJ (version 1.53a). For each section, 8.5 µm-thick Z-stacks of the dorsal CA1 were created using the maximum intensity projection function in ImageJ. For each *st. pyramidale*, *radiatum*, and *lacunosum-moleculare*, two square regions of interest were centered in each stack. Measurements were made from each mouse in each region of interest. The raw integrated density (defined as the sum of the values for all pixels) of the Z-stack region of interest expressing the fluorescent label was measured for the volume of target pixels, and the average of each measurement was taken as representative for the region and each mouse.

### AAV injections

Mice were anesthetized in a closed chamber filled with the inhalation anesthetic isoflurane (RWD Life Science, R510-22-10) and then fixed in a stereotaxic apparatus (Stoelting Co.). Anesthesia was maintained with isoflurane inhalation (1%−2.5% via trachea). The eyes of the mice were safeguarded using erythromycin ophthalmic ointment (0.5%). The skull was exposed and cleaned using 3% hydrogen peroxide. Small holes in the skull were then drilled with the following stereotaxic coordinates: left hippocampus (triple injection: AP: -1/ ML: -0.7/ DV: - 1.65; AP: -1.8/ ML: -1.5/ DV: -2; AP: -2.7/ ML: -2/ DV: -2, below the skull surface) and right hippocampus (triple injection: AP: -1/ ML: +0.7/ DV: -1.65; AP: -1.8/ ML: +1.5/ DV -2; AP: - 2.7/ ML: +2/ DV: -2, below the skull surface). The virus was injected using a 34-gauge needle with a Hamilton syringe at 0.1 µl/min rate into target regions. At all injected points, the tip of the needle was positioned 0.05 mm below the target coordinate and returned to the target site after 2 min. After injection, the needle stayed in place for an additional 7 min and was slowly withdrawn. AAVs expressing Cre-recombinase and eGFP were from Addgene. For physiology, 0.5 µl of virus pENN.AAV.CMVs.PI.Cre.rBG (AAV2.9) (1 x 10^13^ viral genomes [vg]/ml) was injected into one CA1, and 0.5 µl of virus pAAV.CMV.PI.EGFP.WPRE.bGH (AAV2.9) (1 x 10^13^ vg/ml) was injected in the contralateral side. Virus pENN.AAV.CamKIIa 0.4 Cre SV40 (AAV2.9) (1 x 10^13^ vg/ml) was used to express Cre-recombinase by the CaMKIIα promoter.

### Conditioning

Active place avoidance was conducted with a commercial computer-controlled system (Bio-Signal Group, Acton, MA). The mouse was placed on a 40-cm diameter circular arena rotating at 1 rpm. The specialized software, Tracker (Bio-Signal Group, Acton, MA), was used to detect the animal’s position 30 times per second by video tracking from an overhead camera. A clear wall made from polyethylene terephthalate glycol-modified (PET-G) was placed on the arena to prevent the animal from jumping off the elevated arena surface. A 5-pole shock grid was placed on the rotating arena, and the shock was scrambled across the 5 poles when the mouse entered the shock zone. All experiments used the “Room+Arena-“ task variant that challenges the mouse on the rotating arena to avoid a shock zone that was a stationary 60° sector (Pastalkova et al., 2006). Every 33 ms, the software determined the mouse’s position, whether it was in the shock zone, and whether to deliver shock. After the animal enters the shock zone for 500 ms, a constant current foot-shock (60 Hz, 500 ms) was delivered and repeated with an interval of 1500 ms until the mouse left the shock zone. The shock intensity was 0.2 or 0.3 mA, which was the minimum amplitude to elicit flinch or escape responses. The animal was forced to actively avoid the designated shock zone because the arena rotation periodically transported it into the shock area. A pretraining period on the apparatus without shock that was equivalent in time to a training session was provided.

The tracked animal positions with timestamps were analyzed offline (TrackAnalysis, Bio-Signal Group, Acton, MA) to extract several end-point measures. The time to first enter the shock zone estimates ability to avoid shock and was taken as an index of between-session long-term place avoidance memory. Short-term memory was assessed by two measures. First, the times to each entry into the shock zone in the first training trial were compared to the times for each entry into the shock zone with the shock off during the pretraining session. Avoidance behavior is observed as an increase in the amplitude of the times for entering the shock zone. Second, the maximum time without receiving a shock was determined for each session. Short-term memory for avoidance behavior is measured as an increase in the maximum time between shocks in the first training trial, compared to the maximum time between entrances into the shock zone with the shock off during pretraining.

For Figure 5 the training schedule was as follows: 1 day after a 30-min pretraining session, the animals received three 30-min training trials, with an intertrial interval of 1 day. Long-term memory retention was tested the following day without shock. Pre-established exclusion criterion was if cannulae were found to be incorrectly targeted. No mice were excluded.

### Statistics

All experiments were performed with blind procedures except for LTP experiments that involved transfection with AAV-eGFP, as the eGFP could be detected visually in the hippocampal slice by the experimenter. Sample sizes vary for the different experimental approaches (biochemistry, extracellular field potential physiology, and behavior). The hypothesis that PKM ζ is compensated predicts all-or-none effects in the experiments, and this provided a basis for sample size estimates. Power analyses were performed using G*Power Version 3.1.9.7 with α = 0.05 and β = 0.8 and large effect sizes of 1.5–2.0. The effect size estimates were based on prior studies that demonstrated essentially all-or-none effects of PKMζ inhibition on the immunoblot, immunohistochemical, physiological, and behavioral assays used here (Hsieh et al., 2021; Tsokas et al., 2024; Tsokas et al., 2016). Two-population Student *t* tests with Bonferroni corrections were performed to compare protein levels by immunoblot and immunohistochemistry in the PKMζ -cKO and control mice. For LTP experiments the responses to test stimuli were averaged across 5 min for statistical comparisons. Repeated measures ANOVA was used to compare the change in the potentiated response at the time points described. Multi-factor comparisons were performed using mixed-design ANOVA with repeated measures, as appropriate. The degrees of freedom for the critical *t* values of the *t* tests and the *F* values of the ANOVAs are reported as subscripts. *Post-hoc* multiple comparisons were performed by Newman-Keuls tests as appropriate. Statistical significance was accepted at *P* < 0.05. Effect sizes for binary comparisons and one-way ANOVAs are reported as Cohen’s *d* and as *η*^2^_p_ for multi-factor ANOVA effects.

**Figure 1 figure supplement 1.**
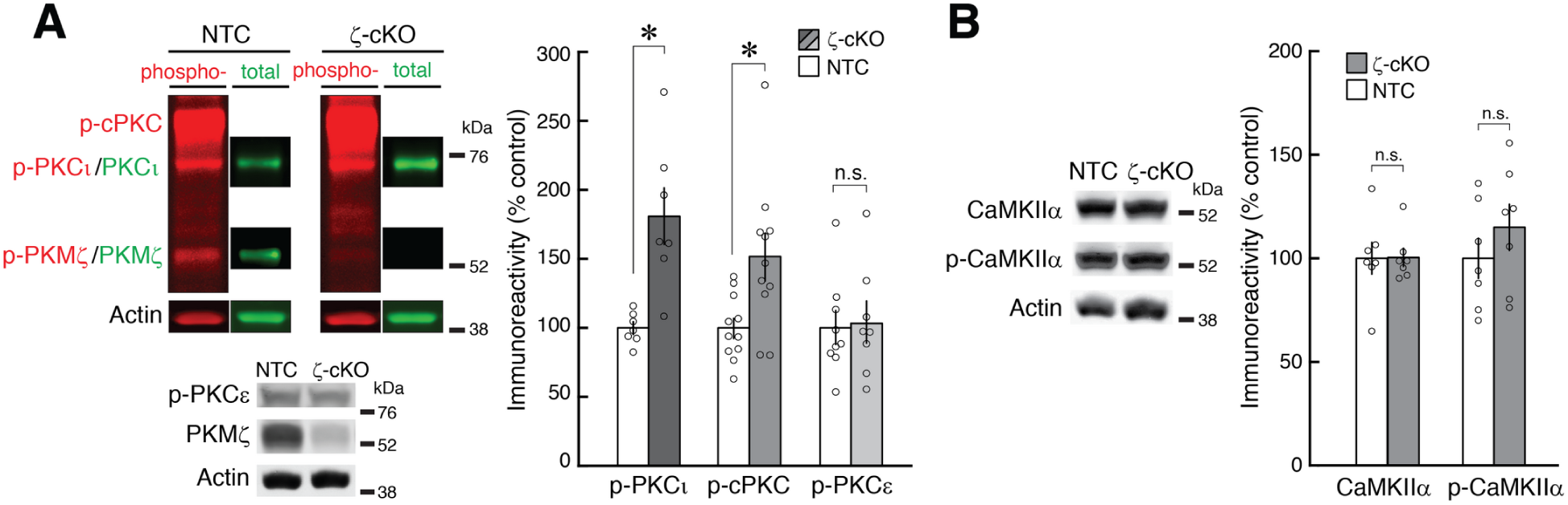
ζ -cKO increases activation-loop-phosphorylation state of atypical PKC ι and conventional PKCs, but not novel PKC ε or T286-autophosphorylation of CaMKIIα. (**A**) Left, above, representative immunoblots show increases in activation-loop phosphorylated PKC ι (p-PKC ι) and conventional-PKCs (p-cPKC) in PKM ζ-cKO mice, compared to non-transgenic controls (NTC). Red, phospho-PKCs; green, total PKCs from the same samples. Mr markers shown at right. Below, p-PKC ε, recognized by its higher Mr, does not change. Right, mean ± SEM (**B**) Levels of total CaMKIIα and T286-autophosphorylated CaMKIIα do not change in ζ-cKO hippocampus. Statistics in Figure 1 — table supplement 2.

**Figure 1 figure supplement 2.**
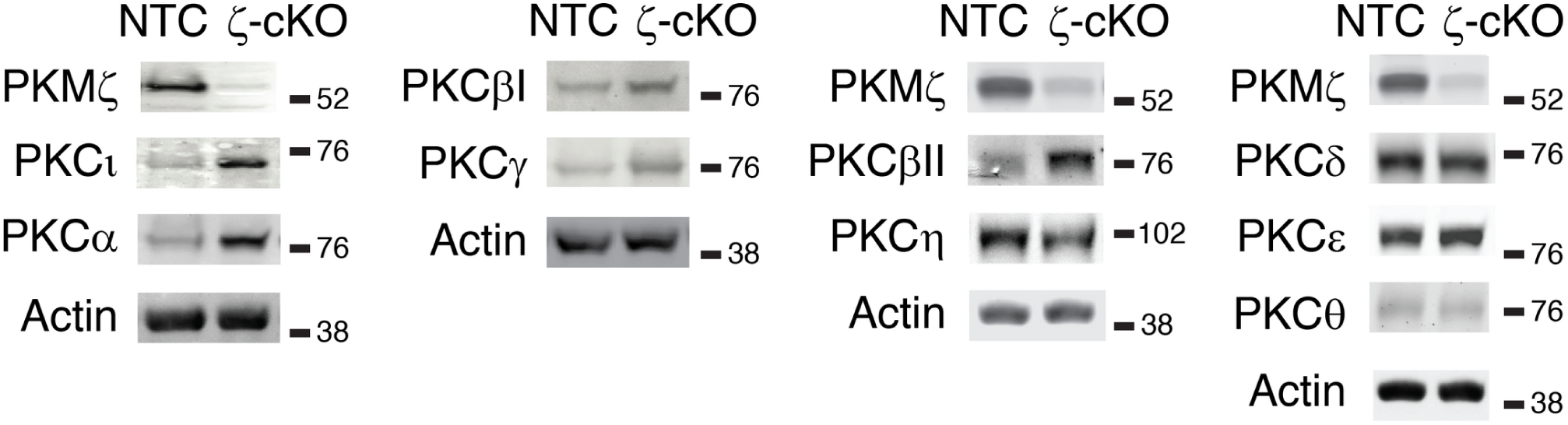
Actin loading controls for immunoblots shown in Figure 1B. The PKM ζ and actin lanes from columns 3 and 4 are from 3 adjacent lanes shown in Figure 1A.

**Figure 4 figure supplement 1.**
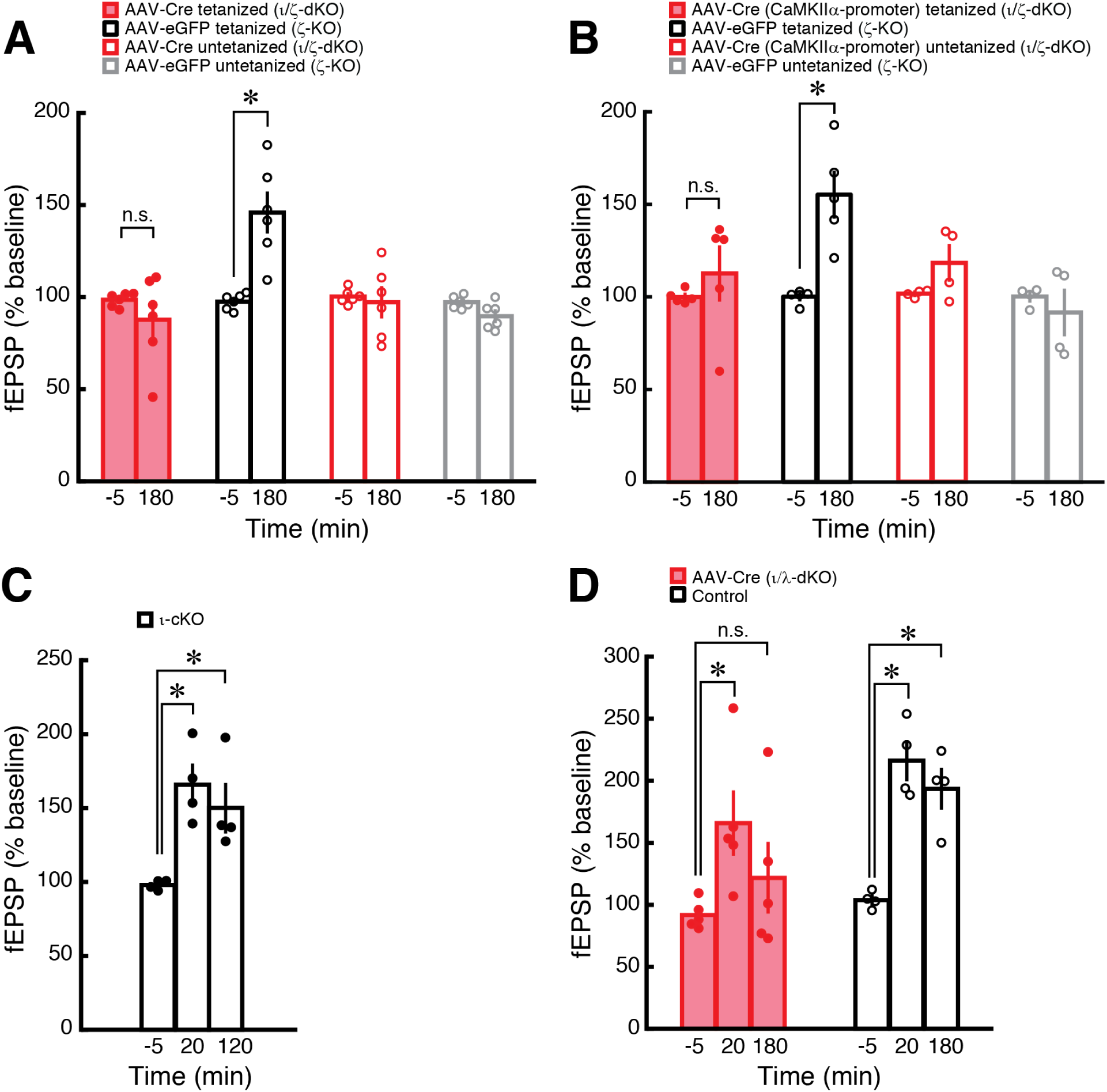
Data used in the statistical analysis of experiments in Figure 4 and Figure 4 — figure supplement 2B. (**A**) Figure 4A, (**B**) Figure 4A, insert, (**C**) Figure 4 — figure supplement 2B, **(D)** Figure 4B.

**Figure 4 figure supplement 2.**
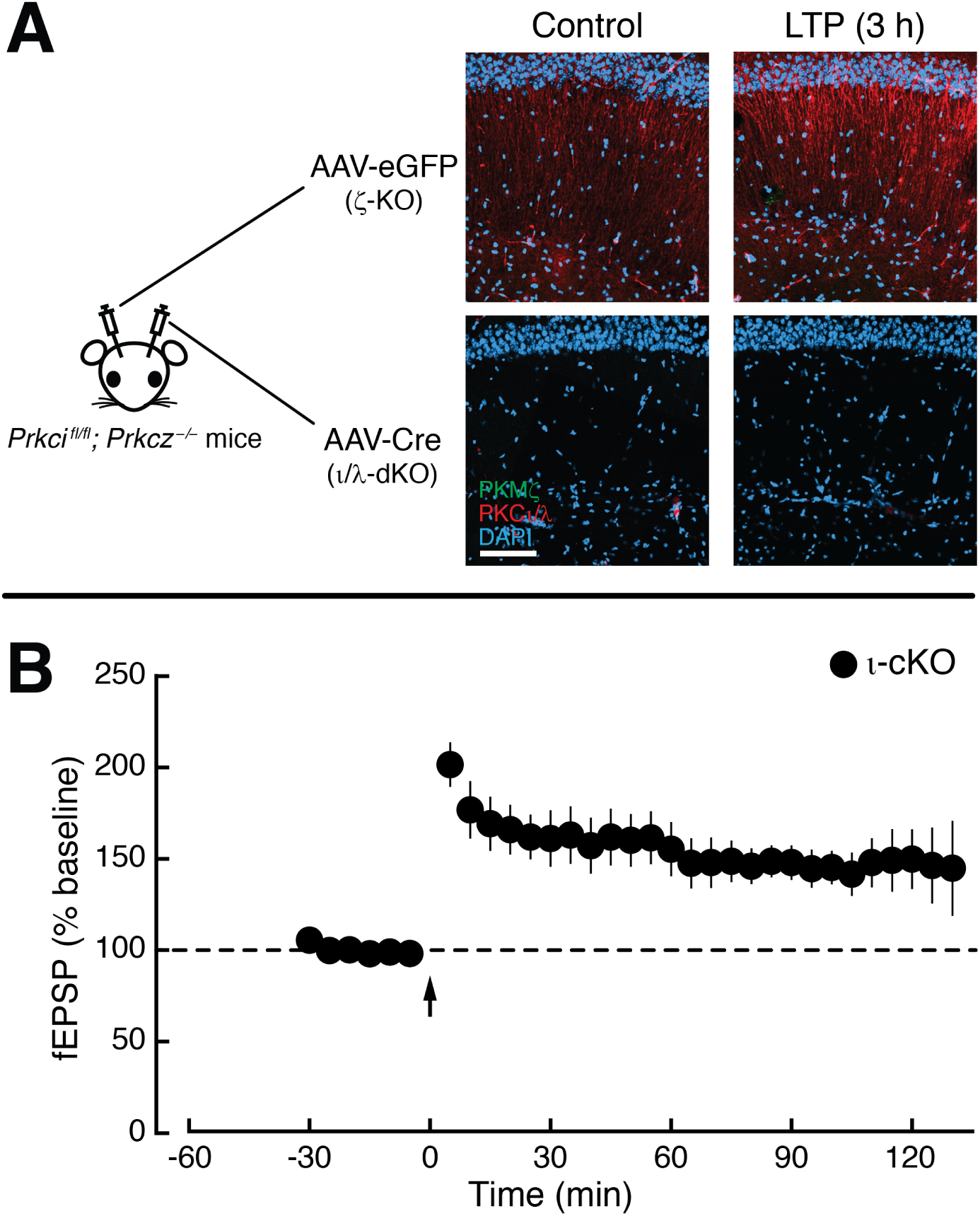
Effects of AAV expression of Cre-recombinase by the CaMKIIα promoter in *Prkci^fl/fl^; Prkcz^−/–^* and *Prkci^fl/fl^; Prkcz^+/+^* mice. (A) Representative immunohistochemistry of AAV expressing Cre-recombinase by CaMKIIα promoter in *Prkci^fl/fl^; Prkcz^−/–^* mice shows loss of PKC ι in ι / ζ-cKO hippocampus and compensatory increase in PKC ι during LTP maintenance in control eGFP-injected hippocampus. Left, schematic of sites of injection; right, PKCι immunohistochemistry. DAPI staining of nuclei shown in blue. Bar = 100 µm. (**B**) ι -cKO (AAV with CaMKIIα promoter expressing Cre-recombinase in hippocampus of *Prkci^fl/fl^; Prkcz^+/+^*mice) shows compensated LTP, as previously described (Sheng et al., 2017). Loss of early-LTP is compensated in the PKC ι -cKO as in prior reports. The repeated measurement ANOVA reveals the main effect of LTP (5-min average of pre-HFS, 20-min, and 2-h post-HFS, *F*_2,6_ = 14.03, *P* = 0.005, *η*^2^_p_ = 0.82) in ι-cKO mice. *Post-hoc* tests confirm that LTP was established at 20 min post-tetanization (5-min pre-HFS vs. 20-min post-HFS, *P* = 0.006) and maintained for 2 h (5-min pre-HFS vs. 120-min post-HFS, *P* = 0.009); n = 4.

**Figure 1 table supplement 1.** Statistics for data presented in (A) Figure 1A and (B) Figure 1B. Significant differences with Bonferroni correction are in bold.

| Isozyme | Non-transgenic control (NTC) | | ζ-cKO | | Degree of freedom | $t$ | $P$ | Cohen's $d$ |
| --- | --- | --- | --- | --- | --- | --- | --- | --- |
| | Mean ± SEM | n | Mean ± SEM | n | | | | |
| (A) |  |  |  |  |  |  |  |  |
| aPKC |  |  |  |  |  |  |  |  |
| PKMζ | 100 ± 4.9 | 9 | 11.8 ± 2.4 | 8 | 15 | 15.4 | < 0.0001 | 7.5 |
| PKCι | 100 ± 7.3 | 4 | 172.4 ± 18.4 | 5 | 7 | 3.9 | 0.01 | 2.2 |
| (B) |  |  |  |  |  |  |  |  |
| cPKC |  |  |  |  |  |  |  |  |
| $\alpha$ | 100 ± 2.3 | 8 | 140.0 ± 7.0 | 8 | 14 | 5.4 | < 0.0001 | 2.7 |
| βI | 100 ± 6.2 | 8 | 142.1 ± 10.6 | 8 | 14 | 3.4 | 0.004 | 1.7 |
| βII | 100 ± 6.3 | 6 | 150.2 ± 5.5 | 5 | 9 | 5.9 | 0.0002 | 3.6 |
| $\gamma$ | 100 ± 8.6 | 9 | 149.6 ± 15.4 | 9 | 16 | 2.8 | 0.01 | 1.3 |
| nPKC |  |  |  |  |  |  |  |  |
| $\delta$ | 100 ± 10.4 | 7 | 108.2 ± 10.3 | 7 | 12 | 0.6 | 0.6 | 0.3 |
| $\varepsilon$ | 100 ± 5.3 | 9 | 114.8 ± 11.0 | 9 | 16 | 1.2 | 0.2 | 0.6 |
| $\eta$ | 100 ± 17.1 | 6 | 109.0 ± 10.9 | 6 | 10 | 0.4 | 0.7 | 0.3 |
| $\theta$ | 100 ± 10.1 | 6 | 97.7 ± 14.9 | 7 | 11 | 0.1 | 0.9 | 0.1 |

**Figure 1 table supplement 2.** Statistics for data presented in (A) Figure 1 — figure supplement 1A and (B) Figure 1 — figure supplement 1B. Significant differences with Bonferroni correction are in bold.

| Kinase | Non-transgenic control (NTC) | | ζ-cKO | | Degree of freedom | $t$ | $P$ | Cohen's $d$ |
| --- | --- | --- | --- | --- | --- | --- | --- | --- |
| | Mean ± S.E.M. | n | Mean ± S.E.M. | n | | | | |
| (A) |  |  |  |  |  |  |  |  |
| p-PKC |  |  |  |  |  |  |  |  |
| p-PKCι | 100 ± 4.0 | 7 | 181.1 ± 19.8 | 7 | 12 | 4.0 | 0.002 | 2.1 |
| p-cPKC | 100 ± 7.1 | 11 | 151.4 ± 16.4 | 11 | 20 | 3.3 | 0.009 | 1.2 |
| p-PKCε | 100 ± 11.7 | 9 | 104.0 ± 14.2 | 8 | 15 | 0.2 | 0.8 | 0.1 |
| (B) |  |  |  |  |  |  |  |  |
| CaMKIIα | 100 ± 7.63 | 7 | 96.33 ± 1.95 | 6 | 11 | 0.43 | 0.67 | 0.24 |
| p-CaMKIIα | 100 ± 9.65 | 7 | 114.94 ± 11.17 | 7 | 12 | 1.01 | 0.33 | 0.54 |

**Figure 2 table supplement 1.** Statistics for data presented in Figure 2B. Significant differences with Bonferroni correction are in bold.

| Isozyme /<br>region | Vehicle |  | OH-TAM |  | Degree of<br>freedom | *t* | *P* | Cohen's *d* |
| --- | --- | --- | --- | --- | --- | --- | --- | --- |
|  | Mean ± SEM | n | Mean ± SEM | n |  |  |  |  |
| PKMζ |  |  |  |  |  |  |  |  |
| *pyramidale* | **100 ± 13.6** | **5** | **14.1 ± 2.1** | **5** | **8** | **6.2** | **0.0003** | **3.9** |
| *radiatum* | **100 ± 3.6** | **5** | **23.6 ± 3.9** | **5** | **8** | **14.4** | **<0.0001** | **9.1** |
| *lac-mol* | 100 ± 22.0 | 5 | 54.2 ± 13.2 | 5 | 8 | 1.8 | 0.1 | 1.1 |
| PKCι |  |  |  |  |  |  |  |  |
| *pyramidale* | **100 ± 9.8** | **5** | **235.3 ± 26.4** | **5** | **8** | **4.8** | **0.001** | **3.0** |
| *radiatum* | **100 ± 36.9** | **5** | **411.2 ± 26.2** | **5** | **8** | **6.9** | **0.0001** | **4.3** |
| *lac-mol* | 100 ± 40.1 | 5 | 226.7 ± 26.7 | 5 | 8 | 2.6 | 0.03 | 1.7 |

## References

Anderson, W. W., & Collingridge, G. L. (2007). Capabilities of the WinLTP data acquisition program extending beyond basic LTP experimental functions. J Neurosci Methods, 162, 346–356. S0165-0270(07)00002-7 [pii] 10.1016/j.jneumeth.2006.12.018

Bailey, C. H., & Kandel, E. R. (1993). Structural changes accompanying memory storage. Annu Rev Physiol, 55, 397–426. 10.1146/annurev.ph.55.030193.002145

Bayer, K. U., & Giese, K. P. (2024). A revised view of the role of CaMKII in learning and memory. Nat Neurosci. 10.1038/s41593-024-01809-x

Chen, S., Cai, D., Pearce, K., Sun, P. Y., Roberts, A. C., & Glanzman, D. L. (2014). Reinstatement of long-term memory following erasure of its behavioral and synaptic expression in Aplysia. Elife, 3, e03896. 10.7554/eLife.03896

Conant, G. C., & Wagner, A. (2004). Duplicate genes and robustness to transient gene knock-downs in Caenorhabditis elegans. Proc Biol Sci, 271, 89–96. 10.1098/rspb.2003.2560

Crick, F. (1984). Memory and molecular turnover. Nature, 312, 101. 10.1038/312101a0

El Allam, A., Alberto, E. J., Mercau, M. E., Dimitrius, T., Pramio, D. T., Krishna, M., Bhat, K. M., Philbrick, W. M., Schechtman, D., Rothlin, C. V., & Ghosh, S. (2024). Functional roles of neural aPKCs in mouse brain development and survival. bioRxiv:2024.05.22.595312. doi: 10.1101/2024.05.22.595312

El-Brolosy, M. A., Kontarakis, Z., Rossi, A., Kuenne, C., Gunther, S., Fukuda, N., Kikhi, K., Boezio, G. L. M., Takacs, C. M., Lai, S. L., Fukuda, R., Gerri, C., Giraldez, A. J., & Stainier, D. Y. R. (2019). Genetic compensation triggered by mutant mRNA degradation. Nature, 568, 193–197. 10.1038/s41586-019-1064-z

El-Brolosy, M. A., & Stainier, D. Y. R. (2017). Genetic compensation: A phenomenon in search of mechanisms. PLoS Genet, 13, e1006780. 10.1371/journal.pgen.1006780

Gu, Z., Steinmetz, L. M., Gu, X., Scharfe, C., Davis, R. W., & Li, W. H. (2003). Role of duplicate genes in genetic robustness against null mutations. Nature, 421, 63–66. 10.1038/nature01198

Han, J., Grau-Perales, A., Harris, R. M., Kao, H.-Y., Pal, A., Alarcon, J. M., Sacktor, T. C., Martiniani, S., Hofmann, H. A., & Fenton, A. A. (2026). Persistently increased expression of PKMzeta and unbiased gene expression profiles identify hippocampal molecular traces of a long-term active place avoidance memory and ‘shadow’ proteins. Adv Science, 0:e21254. 10.1002/advs.202521254

Hebb, D. O. (1949). The Organization of Behavior. A Neuropsychological Theory. New York John Wiley and Sons, Inc. London.

Hernandez, A. I., Blace, N., Crary, J. F., Serrano, P. A., Leitges, M., Libien, J. M., Weinstein, G., Tcherapanov, A., & Sacktor, T. C. (2003). Protein kinase Mζ synthesis from a brain mRNA encoding an independent protein kinase Cζ catalytic domain. Implications for the molecular mechanism of memory. J Biol Chem, 278, 40305–40316. 10.1074/jbc.M307065200

Hsieh, C., Cano, D. A., Tsokas, P., Cottrell, J. E., Fenton, A. A., Shouval, H., & Sacktor, T. C. (2026). PKMzeta-KIBRA interactions, molecular turnover, and memory. Mol Brain (in press). 10.64898/2026.01.05.697418

Hsieh, C., Tsokas, P., Grau-Perales, A., Lesburgueres, E., Bukai, J., Khanna, K., Chorny, J., Chung, A., Jou, C., Burghardt, N. S., Denny, C. A., Flores-Obando, R. E., Hartley, B. R., Rodriguez Valencia, L. M., Hernandez, A. I., Bergold, P. J., Cottrell, J. E., Alarcon, J. M., Fenton, A. A., & Sacktor, T. C. (2021). Persistent increases of PKMzeta in memory-activated neurons trace LTP maintenance during spatial long-term memory storage. Eur J Neurosci, 54, 6795–6814. 10.1111/ejn.15137

Jiang, J., Parameshwaran, K., Seibenhener, M. L., Kang, M. G., Suppiramaniam, V., Huganir, R. L., Diaz-Meco, M. T., & Wooten, M. W. (2009). AMPA receptor trafficking and synaptic plasticity require SQSTM1/p62. Hippocampus, 19, 392–406. 10.1002/hipo.20528

Kniffin, A. R., English, E. A., & Briand, L. A. (2025). PKMzeta is necessary for long-term depression and long-term potentiation in the medial prefrontal cortex. J Physiol. 10.1113/JP289373

Lee, A. M., Kanter, B. R., Wang, D., Lim, J. P., Zou, M. E., Qiu, C., McMahon, T., Dadgar, J., Fischbach-Weiss, S. C., & Messing, R. O. (2013). Prkcz null mice show normal learning and memory. Nature, 493, 416–419. 10.1038/nature11803

Lisman, J. (2017). Criteria for identifying the molecular basis of the engram (CaMKII, PKMzeta). Mol Brain, 10, 55. 10.1186/s13041-017-0337-4

Lisman, J. E. (1985). A mechanism for memory storage insensitive to molecular turnover: a bistable autophosphorylating kinase. Proc Natl Acad Sci U S A, 82, 3055–3057. 10.1073/pnas.82.9.3055

Ma, Z., Zhu, P., Shi, H., Guo, L., Zhang, Q., Chen, Y., Chen, S., Zhang, Z., Peng, J., & Chen, J. (2019). PTC-bearing mRNA elicits a genetic compensation response via Upf3a and COMPASS components. Nature, 568, 259–263. 10.1038/s41586-019-1057-y

Matus, A. (2000). Actin-based plasticity in dendritic spines. Science, 290, 754–758. http://www.ncbi.nlm.nih.gov/entrez/query.fcgi?cmd=Retrieve&db=PubMed&dopt=Citati on&list_uids=11052932

Mercau, M. E., Hackbarth, R.-M., Liu, X., Wang, H., Zhang, L., Basu, M. K., Rothlin, C. V., & Ghosh, S. (2024). Time-resolved function of cell polarity kinases PRKCZ and PRKCI in CNS myelination. bioRxiv:2024.04.22.589759.

Miller, S. G., & Kennedy, M. (1986). Regulation of brain type II Ca2+/calmodulin-dependent protein kinase by autophosphorylation: a Ca2+-triggered molecular switch. Cell, 44, 861–870.

O’Brian, C. A., Liskamp, R. M., Solomon, D. H., & Weinstein, I. B. (1985). Inhibition of protein kinase C by tamoxifen. Cancer Res, 45, 2462–2465. https://www.ncbi.nlm.nih.gov/pubmed/3157445

Parker, S. S., Mandell, E. K., Hapak, S. M., Maskaykina, I. Y., Kusne, Y., Kim, J. Y., Moy, J. K., St John, P. A., Wilson, J. M., Gothard, K. M., Price, T. J., & Ghosh, S. (2013). Competing molecular interactions of aPKC isoforms regulate neuronal polarity. Proc Natl Acad Sci U S A, 110, 14450–14455. 10.1073/pnas.1301588110

Pastalkova, E., Serrano, P., Pinkhasova, D., Wallace, E., Fenton, A. A., & Sacktor, T. C. (2006). Storage of spatial information by the maintenance mechanism of LTP. Science, 313, 1141–1144. 10.1126/science.1128657

Ramón y Cajal, S. (1894). La fine structure des centres nerveux. 55, 444–468 (1894). Proc. R. Soc. Lond., 55, 444–468.

Ren, S. Q., Yan, J. Z., Zhang, X. Y., Bu, Y. F., Pan, W. W., Yao, W., Tian, T., & Lu, W. (2013). PKClambda is critical in AMPA receptor phosphorylation and synaptic incorporation during LTP. EMBO J, 32, 1365–1380. 10.1038/emboj.2013.60

Sacktor, T. C. (2026). PKMzeta-knockout mice lack neocortical long-term potentiation: limits of hippocampal compensation and differential memory rescue. J Physiol, 604, 1009–1010. 10.1113/JP290246

Sacktor, T. C., & Fenton, A. A. (2018). What does LTP tell us about the roles of CaMKII and PKMzeta in memory? Mol Brain, 11, 77. 10.1186/s13041-018-0420-5

Sacktor, T. C., Osten, P., Valsamis, H., Jiang, X., Naik, M. U., & Sublette, E. (1993). Persistent activation of the zeta isoform of protein kinase C in the maintenance of long-term potentiation. Proc.Natl.Acad.Sci.USA, 90, 8342–8346. (In File)

Scharf, M. T., Woo, N. H., Lattal, K. M., Young, J. Z., Nguyen, P. V., & Abel, T. (2002). Protein synthesis is required for the enhancement of long-term potentiation and long-term memory by spaced training. J Neurophysiol, 87, 2770–2777. http://www.ncbi.nlm.nih.gov/entrez/query.fcgi?cmd=Retrieve&db=PubMed&dopt=Citati on&list_uids=12037179

Schwartz, J. H. (1993). Cognitive kinases. Proc Natl Acad Sci U S A, 90, 8310–8313. http://www.ncbi.nlm.nih.gov/entrez/query.fcgi?cmd=Retrieve&db=PubMed&dopt=Citati on&list_uids=8104334

Scott, J., Thakar, S., Mao, Y., Qin, H., Hejran, H., Lee, S. Y., Yu, T., Klezovitch, O., Cheng, H., Mu, Y., Ghosh, S., Vasioukhin, V., & Zou, Y. (2019). Apical-Basal Polarity Signaling Components, Lgl1 and aPKCs, Control Glutamatergic Synapse Number and Function. iScience, 20, 25–41. 10.1016/j.isci.2019.09.005

Seidl, S., Braun, U., Roos, N., Li, S., Ludtke, T. H., Kispert, A., & Leitges, M. (2013). Phenotypical analysis of atypical PKCs in vivo function display a compensatory system at mouse embryonic day 7.5. PLoS One, 8, e62756. 10.1371/journal.pone.0062756

Serrano, P., Yao, Y., & Sacktor, T. C. (2005). Persistent phosphorylation by protein kinase Mζ maintains late-phase long-term potentiation. J Neurosci, 25, 1979–1984. 10.1523/JNEUROSCI.5132-04.2005

Shema, R., Haramati, S., Ron, S., Hazvi, S., Chen, A., Sacktor, T. C., & Dudai, Y. (2011). Enhancement of consolidated long-term memory by overexpression of protein kinase Mzeta in the neocortex. Science, 331, 1207–1210. 331/6021/1207 [pii] 10.1126/science.1200215

Shema, R., Sacktor, T. C., & Dudai, Y. (2007). Rapid erasure of long-term memory associations in the cortex by an inhibitor of PKMzeta. Science, 317, 951–953. 317/5840/951 [pii] 10.1126/science.1144334

Sheng, T., Wang, S., Qian, D., Gao, J., Ohno, S., & Lu, W. (2017). Learning-Induced Suboptimal Compensation for PKCiota/lambda Function in Mutant Mice. Cereb Cortex, 27, 3284–3293. 10.1093/cercor/bhx077

Shouval, H. Z., Hsieh, C., Flores-Obando, R. E., Cano, D. A., Tracy, T. E., & Sacktor, T. C. (2025). Maintenance of memory by negative feedback of synaptic protein elimination: modeling KIBRA-PKMzeta dynamics in LTP. Learn Mem, 32. 10.1101/lm.054077.124

Tsien, R. Y. (2013). Very long-term memories may be stored in the pattern of holes in the perineuronal net. Proc Natl Acad Sci U S A, 110, 12456–12461. 10.1073/pnas.1310158110

Tsokas, P., Grace, E. A., Chan, P., Ma, T., Sealfon, S. C., Iyengar, R., Landau, E. M., & Blitzer, R. D. (2005). Local protein synthesis mediates a rapid increase in dendritic elongation factor 1A after induction of late long-term potentiation. J Neurosci, 25, 5833–5843. 10.1523/JNEUROSCI.0599-05.2005

Tsokas, P., Hsieh, C., Flores-Obando, R. E., Bernabo, M., Tcherepanov, A., Hernandez, A. I., Thomas, C., Bergold, P. J., Cottrell, J. E., Kremerskothen, J., Shouval, H. Z., Nader, K., Fenton, A. A., & Sacktor, T. C. (2024). KIBRA anchoring the action of PKMzeta maintains the persistence of memory. Sci Adv, 10, eadl0030. 10.1126/sciadv.adl0030

Tsokas, P., Hsieh, C., Yao, Y., Lesburgueres, E., Wallace, E. J., Tcherepanov, A., Jothianandan, D., Hartley, B. R., Pan, L., Rivard, B., Farese, R. V., Sajan, M. P., Bergold, P. J., Hernandez, A. I., Cottrell, J. E., Shouval, H. Z., Fenton, A. A., & Sacktor, T. C. (2016). Compensation for PKMzeta in long-term potentiation and spatial long-term memory in mutant mice. Elife, 5, e14846. 10.7554/eLife.14846

Tsokas, P., Ma, T., Iyengar, R., Landau, E. M., & Blitzer, R. D. (2007). Mitogen-activated protein kinase upregulates the dendritic translation machinery in long-term potentiation by controlling the mammalian target of rapamycin pathway. J Neurosci, 27, 5885–5894. https://doi.org/27/22/5885 [pii] 10.1523/JNEUROSCI.4548-06.2007

Tsokas, P., Rivard, B., Hsieh, C., Cottrell, J. E., Fenton, A. A., & Sacktor, T. C. (2019). Antisense Oligodeoxynucleotide Perfusion Blocks Gene Expression of Synaptic Plasticity-related Proteins without Inducing Compensation in Hippocampal Slices. Bio Protoc, 9. 10.21769/BioProtoc.3387

Tullis, J. E., Larsen, M. E., Rumian, N. L., Freund, R. K., Boxer, E. E., Brown, C. N., Coultrap, S. J., Schulman, H., Aoto, J., Dell’Acqua, M. L., & Bayer, K. U. (2023). LTP induction by structural rather than enzymatic functions of CaMKII. Nature, 621, 146–153. 10.1038/s41586-023-06465-y

Volk, L. J., Bachman, J. L., Johnson, R., Yu, Y., & Huganir, R. L. (2013). PKM-zeta is not required for hippocampal synaptic plasticity, learning and memory. Nature, 493, 420–423. 10.1038/nature11802

Wang, S., Sheng, T., Ren, S., Tian, T., & Lu, W. (2016). Distinct Roles of PKCiota/lambda and PKMzeta in the Initiation and Maintenance of Hippocampal Long-Term Potentiation and Memory. Cell Rep, 16, 1954–1961. 10.1016/j.celrep.2016.07.030

White, J. K., Gerdin, A. K., Karp, N. A., Ryder, E., Buljan, M., Bussell, J. N., Salisbury, J., Clare, S., Ingham, N. J., Podrini, C., Houghton, R., Estabel, J., Bottomley, J. R., Melvin, D. G., Sunter, D., Adams, N. C., Sanger Institute Mouse Genetics, P., Tannahill, D., Logan, D. W., … Steel, K. P. (2013). Genome-wide generation and systematic phenotyping of knockout mice reveals new roles for many genes. Cell, 154, 452–464. 10.1016/j.cell.2013.06.022

Zhang, L., & Wei, X. (2022). The Roles of Par3, Par6, and aPKC Polarity Proteins in Normal Neurodevelopment and in Neurodegenerative and Neuropsychiatric Disorders. J Neurosci, 42, 4774–4793. 10.1523/JNEUROSCI.0059-22.2022

